# Seminal fluid proteins can mitigate sexual conflict: the case of remating in insects

**DOI:** 10.64898/2026.02.27.708467

**Authors:** Piotr Michalak, David Duneau, Jean-Baptiste Ferdy

## Abstract

Males during mating never transfer just sperm; to the best of our knowledge, they always deliver a rich seminal fluid as well. The proteins in the seminal fluid have an important sperm supporting role, but they also cause changes in the female physiology and can impose a mating cost. The associated costs and delay in the time of remating, lead to the view that those proteins evolved primarily due to sexual conflict and delay female remating beyond the optimal rate. To examine the role of seminal fluid proteins in sexual conflict we use a mathematical model of reproductive physiology, informed by the accumulated knowledge on *Drosophila melanogaster*. In accordance with the theory, we find that males always benefit from inducing longer remating intervals in females. But, we also find that this conflict is reduced when female reproduction is regulated by the male proteins. Without seminal fluid proteins females have a single, well-defined, optimal remating rate. However, when seminal proteins are used to regulate females reproduction, females can reach the same offspring production for a range of mating intervals. This wider range of possible remating times could provide females with a buffer against uncertain mating opportunity. It could also allow females to be more selective on male quality, by reducing the cost associated with delaying remating. Our results suggest that, while there is a conflict over the remating rate, seminal fluid proteins reduce its intensity, highlighting their role in aligning the interests of both sexes.

## 1 Introduction

When fertilizing eggs internally, males deliver a seminal fluid containing both sperm and a range of accessory proteins. These seminal fluid proteins play key roles in supporting sperm and are known to nourish sperm cells, enhance their motility, and survival within the female reproductive tract (reviewed in 1–5). In addition to supporting sperm function, seminal proteins have a range of effects on female physiology (1, 2, 6–10). Those proteins stimulate ovulation in camels, promote embryo growth in mice before implantation (11–14) and prepare the female immune system to tolerate sperm and embryo in mammals (e.g. 15). In insects, seminal fluid proteins stimulate egg production, ovulation, sperm storage and release—but can also lead to increased vulnerability to infection and change female behavior (9, 10, 16–24). Because seminal fluid proteins are produced by males and modify females physiology or behavior after mating (2, 3, 25–29), they have often been considered as a way for males to exert control over female reproduction. However, the extent to which seminal proteins are key actors in the sexual conflict remains an open question which needs to be addresses to better understand the evolution of reproductive interactions.

Female remating behavior is a key area where male and female reproductive interests may diverge. In species where females store sperm to continuously fertilize eggs after mating, female remating is generally detrimental to the previous mate. This is because the stored sperm will either be displaced by subsequent males’ sperm, or at least will not sire all future offspring (30). As a result, males are expected to evolve strategies delaying female remating for as long as possible. Most females exhibit a refractory period after mating during which they do not mate. This period, here referred to as the remating interval, is partly influenced by the presence of various seminal fluid proteins. In insects, mice, nematodes, and some primates, seminal proteins play a key role in forming of a mating plug, which can physically block the female reproductive tract, preventing remating (5, 31, 32). In other animals, including rats, bumblebees, grasshoppers, moths, snails, and fruit flies, these proteins modify female behavior and delay remating by reducing female receptivity or reducing female attractiveness to males (3, 17, 32–35). Irrespective of the mechanism at play, the extension of the female refractory period is often interpreted as being in male’s interest and therefore a clear example of male-driven manipulation mediated by seminal fluid proteins.

However, the same observations can be interpreted from a female perspective. Females could delay remating to optimize their own reproduction and use seminal proteins as cues to adjust their reproductive physiology (2, 3, 25–27). Here we will investigate this possibility in the case of sperm-storing insects. In these species, reproductive output declines as stored sperm becomes depleted. Females who continue to produce and lay eggs beyond sperm depletion would waste resources by laying unfertilized eggs. Efficient reproduction involves therefore need to adjust egg production to match sperm availability, and remate when sperm stores are too low. The decision to remate involves balancing the benefits of replenishing sperm storage—maintaining high reproductive output and potentially increasing offspring genetic diversity (36, 37)—against the potential costs of additional matings. Critically, both adjusting the egg production and decision to remate require that females can monitor the number of stored sperm (3, 25, 38). Our recent theoretical work suggests that seminal fluid proteins can serve as reliable indicators of sperm availability (38).

This theory has some support in *Drosophila melanogaster*, where a seminal fluid protein called *sex peptide* (SP) delays remating. SP decreases female lifespan and has for this reason long been interpreted as acting in males interest only (39, 40). But it has been recently suggested that SP could optimize fertilization by coordinating the release of eggs and that of sperm (3, 25) (although this coordination is not perfect (41)). SP might be therefore beneficial to females and may be used by both sexes to pursue their respective strategies regarding the optimization of remating intervals.

Studying of the role of SP in sexual conflicts over remating involves comparing the optimal remating strategies of females and males. Measuring differences between females and males strategies while controlling for the various other effects of SP would be highly complex, if even possible, in experimental systems. But a mathematical model can help testing the idea, provided that it does include a good description of female reproductive physiology. There exists a large body of literature on the effects of SP in *Drosophila* (reviewed in 3). This provides an invaluable resource to build a mathematical model that adequately reflects the physiology of *D. melanogaster*. We have previously proposed such a model (38), which showed that females using seminal fluid proteins to coordinate the release of sperm and eggs are better able to optimize their reproduction. We adapted the model here to test the hypothesis that the use of these proteins also affects the optimal interval between matings. To do so, we included multiple matings in our model, considering that females reproductive output is limited by both their lifespan and their resource budget. We then compared the optimal remating interval (i.e. the strategy resulting in the highest number of offspring) under two scenarios: a realistic scenario where females use seminal fluid to regulate their reproductive physiology—referred to as SRP for “Seminal fluid proteins Regulated Physiology”, and a hypothetical scenario where females lack this ability (CR, “Constant Reproduction”). Finally, we compared the optimal female and male remating strategies in both scenarios to determine if seminal fluid proteins, chiefly SP, influence the intensity of the conflict. We found that the optimized reproduction under SRP reduces females sensitivity to delayed remating, and consequently lowers the intensity of the intersexual conflict.

## 2 Model description

The model uses a set of differential equations to predict the three quantities which directly determine how many offspring a female produces at any given time *t*: the number of produced eggs, the number of sperm released from storage, and the number of eggs released (oviposited) (Fig. 1A). We considered two scenarios: one in which there is no regulation because the parameters that control the dynamics of these quantities are fixed (CR), and one where they are regulated by seminal fluid proteins (SRP).

**Figure 1:**
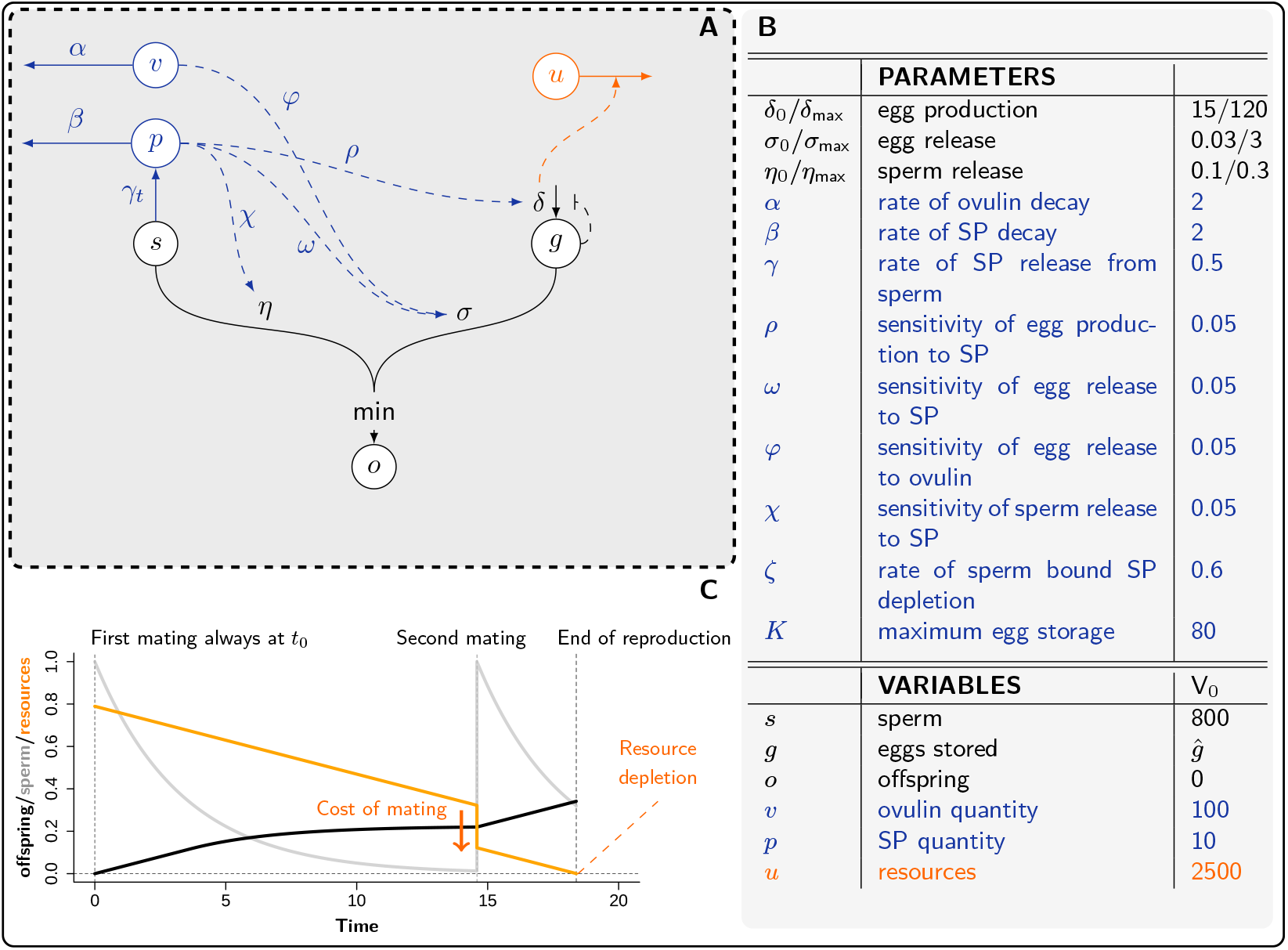
Model of reproduction regulated by seminal fluid proteins based on (38) extended for multiple matings. (A) Schematic representation of the model. The produced offspring (*o*) equals the number of released sperm (*ηs*) or of released eggs (*σg*), depending on which one is limiting. The number of viable eggs in storage (*g*) is determined by egg production (*δ*) and egg release (*σ*). The number of sperm (*s*) is determined by the number stored upon mating (*s*_0_) and by sperm release (*η*). Without regulation production and release parameters are constant; with regulation they are modulated by the seminal fluid proteins (dashed blue arrows). Ovulin (*v*) and sex peptides (SP, *p*) follow independent dynamics (solid blue arrows), with SP being gradually released from sperm (*γ*). Both egg production and mating incur costs, which depletes the resource budget (*u*). (B) Parameters’ values and initial state of the variables. When feasible, values have been chosen so as to match data available for *D. melanogaster* (see 38). The values given are default for the simulations, except when otherwise indicated. Parameters with 0 subscript correspond to base values, while those with *max* subscript are the maxima. (C) Example dynamics when the number of sperm is used as a signal to trigger remating. At first mating, sperm (gray line) are stored and deplete at constant rate. Offspring are produced continuously after mating, limited by either sperm or eggs availability. Females remate when the number of sperm falls below certain threshold. Sperm are then replenished and offspring production restarts. The resources (orange) are expended on egg production or to pay a fixed cost of mating (here *cm* = 500). Female reproduction stops when resources reach zero, or reach the maximum lifespan (100 days). To allow for representation on shared axis, all values were normalized so that maximal value is one.

The model with no regulation, which is similar to that of Gilbert et al. (42), is determined by the three following differential equations, with *g* the number of eggs in the ovarioles, *s* the number of sperm stored in spermathecae or seminal receptacle and *o* the number of offspring produced since the time of mating.

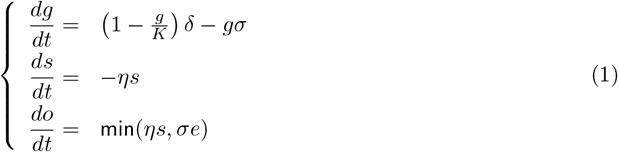

In the first equation, which predicts how many unfertilized eggs are stored, *K* is the maximum egg storage capacity, *δ* the number of eggs produced per unit of time and *σ* the rate at which eggs are released. The second equation describes how sperm are released from storage over time and *η* is the rate of sperm release. The third equation describes the production of offspring. We assume that the number of offspring produced per unit time is set by whichever gamete (sperm or egg) is limiting at time *t*. This implies that if sperm release exceeds that of eggs, sperm are being wasted. In the reverse situation, the female would waste eggs by releasing some that are not fertilized.

In (1), the parameters controlling female physiology are constant—seminal proteins have no influence on physiology. This hypothetical case of constant reproduction model (CR) does not represent a naturally occurring case, nor any experimental mutant. Rather it is a hypothetical scenario where females had evolved without any cues to regulate her physiology. We will use this scenario as a reference to determine the importance of such cues.

A scenario where females use seminal proteins as cues to regulate their reproductive physiology (SRP) is more realistic. It requires that the parameters determining egg production (*δ*), egg release (*σ*) and sperm release (*η*) are modulated by ovulin-like (*v*) and sex peptide-like (SP, *p*) proteins. This modulation is described by the following set of equations

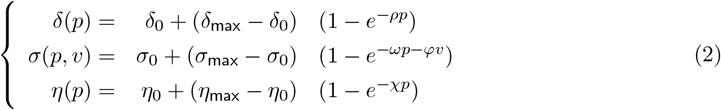

where *ρ, ω, φ* and *χ* can be interpreted as sensitivity parameters (for egg production, egg release and sperm release, respectively) to changes in proteins concentration (*p* and *v*). Parameters with 0 subscript are base values for the three parameters—rates that are achieved in the complete absence of seminal proteins. Parameters with max subscript are the maximum values of the same parameters, reached at high protein concentrations. The degree of seminal proteins influence on the three processes is therefore determined by the difference between these base and maximal values. For example, if the maximum values of parameters equal the base values, seminal fluid proteins have no influence, as in the CR scenario. Similarly if the sensitivity parameters (*ρ, ω, φ* and *χ*) are zero.

The two proteins we considered in our model are provided to females during mating. Their dynamics are described by

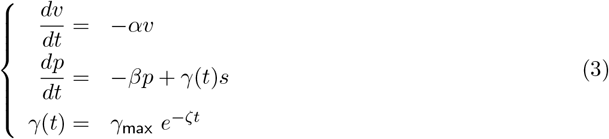

We considered that ovulin is transferred during mating and then simply decays over time at a constant rate *α*. SP (sex-peptide) dynamics are more complex, because SP is attached to sperm and gradually released into the female reproductive tract after mating. SP thus decay after mating at rate *β*, but are also released and supplied from sperm at rate *γ*. SP from sperm depletes over time, because the stock of SP attached to sperm is set at the time of mating. We therefore made *γ* a decreasing function of time which approaches zero when *t* gets large.

We parameterize the model so that a unit of time corresponds to approximately one day. We also aimed to set parameter values so that they match the processes described in the literature (see Fig. 1B, and (38) for justification). Our previous analysis of this model focused on a single mating and suggested that SP can be an efficient cue to monitor the number of stored sperm. SRP model indeed allows to align sperm and egg release, so that egg wasting is limited (38), which benefits both males and females.

Our main goal here is to investigate how regulation using seminal proteins influences females’ remating decisions. For this purpose, we first modeled multiple mating as an iteration of the single mating model. Each time a female remates, she replenishes her stock of sperm and obtains new batch of seminal fluid proteins. Formally, this is expressed as

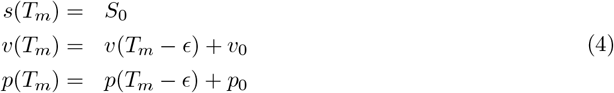

where *T*_*m*_ is the time at which the female mates again, and *Tm*−*ϵ* the time just before remating. At remating, sperm in storage (*s*) is set to maximum number a female can store. This means females can never exceed her sperm storage capacity (*S*_0_). However, the newly provided free proteins (*v*_0_ and *p*_0_) add to those available before remating. We also assumed that the stock of SP attached to sperm is replenished upon remating. This means that the depletion of SP from sperm is based on the time since the last mating, and all sperm are treated as identical. Because SP is replenished at mating, the value of *γ*(*t*) is now given by

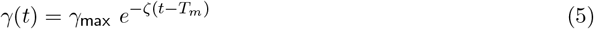

with *t > T*_*m*_.

The offspring production dynamics always begin with mating (at *t* = 0, Fig. 1C). Subsequent matings occur when specific stimulus is reached. We consider three mutually exclusive scenarios determining female remating decisions. The first and simplest is based on time: in this scenario, female waits a fixed time interval before she remates. This is similar to seasonal or cyclical reproduction. The second and third scenarios are based on the state of variable corresponding to physiological characteristic. The decision to remate can be initiated either when sperm left in storage (*s*) fall below a fixed threshold (second scenario) or when the quantity of SP (*p*) does (third scenario).

A model of multiple matings has to account for the fact that females cannot mate an infinite number of times. We assumed that females have a maximal lifespan set to 100 time units (approximately 100 days). Because egg production, and possibly mating, are costly, females are assigned a fixed amount of resources they can expend on those functions. We use resources in a broad sense, encompassing both energy and other resources available to the female during her lifetime. The resource dynamics are described by:

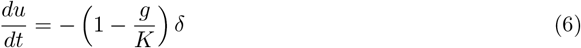

where *u* is the remaining resource budget. We have assumed that one unit of resources is spent for each new egg produced, so that the resources spent are directly proportional to the number of eggs produced. We fixed the initial resource budget to *U*_0_ = 2500, which means that females have resources to produce a maximum of 2500 eggs in their reproductive lifespan, which is within, but high, physiological upper limits (43–45). We also stipulated that mating is costly. Such costs could come either from male harassment, or from transmission of harmful pathogens, which expose females to infections and lead to survival trade-offs of mounting an immune response. (18, 46). We modeled this by subtracting a fixed cost of mating (*cm*) from the resources remaining at the time of mating (*T*_*m*_):

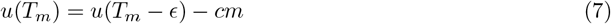

In the model, a female will stop reproducing when she reaches her lifespan, or when she has used up all her resources. If the stimulus triggering remating is reached, but the cost of mating is higher than the resources left, the female does not remate.

## 3 Seminal fluid proteins preserve fitness when time between matings is long

To understand how Seminal fluid proteins Regulated Physiology (SRP) influences the female optimal remating, we first compared females’ reproductive dynamics under Constant Reproduction (CR model) and SRP scenarios. We examined the resulting dynamics under the three proposed remating stimuli—time, sperm and SP. After first mating (*t* = 0, Fig. 2A), the female begins reproduction. Under CR, offspring production is initially constant, because eggs are limiting and produced at a constant rate. Offspring production starts to slow down when there are not enough sperm left to fertilize all the eggs the female lays. When a second mating occurs, sperm supplies are replenished and offspring production returns to its maximal value. This pattern repeats until the female even-tually runs out of resources or dies. This cyclical time pattern remains unchanged when the number of sperm or the quantity of SP are the stimulus which initiate remating. In fact, threshold values for sperm and SP can be set so that the cycles are perfectly identical in the three scenarios (Figures 2A-C). Thus, under CR, the time and SP stimuli directly depend on time and yield the same dynamics of reproduction as when remating time interval is fixed.

**Figure 2:**
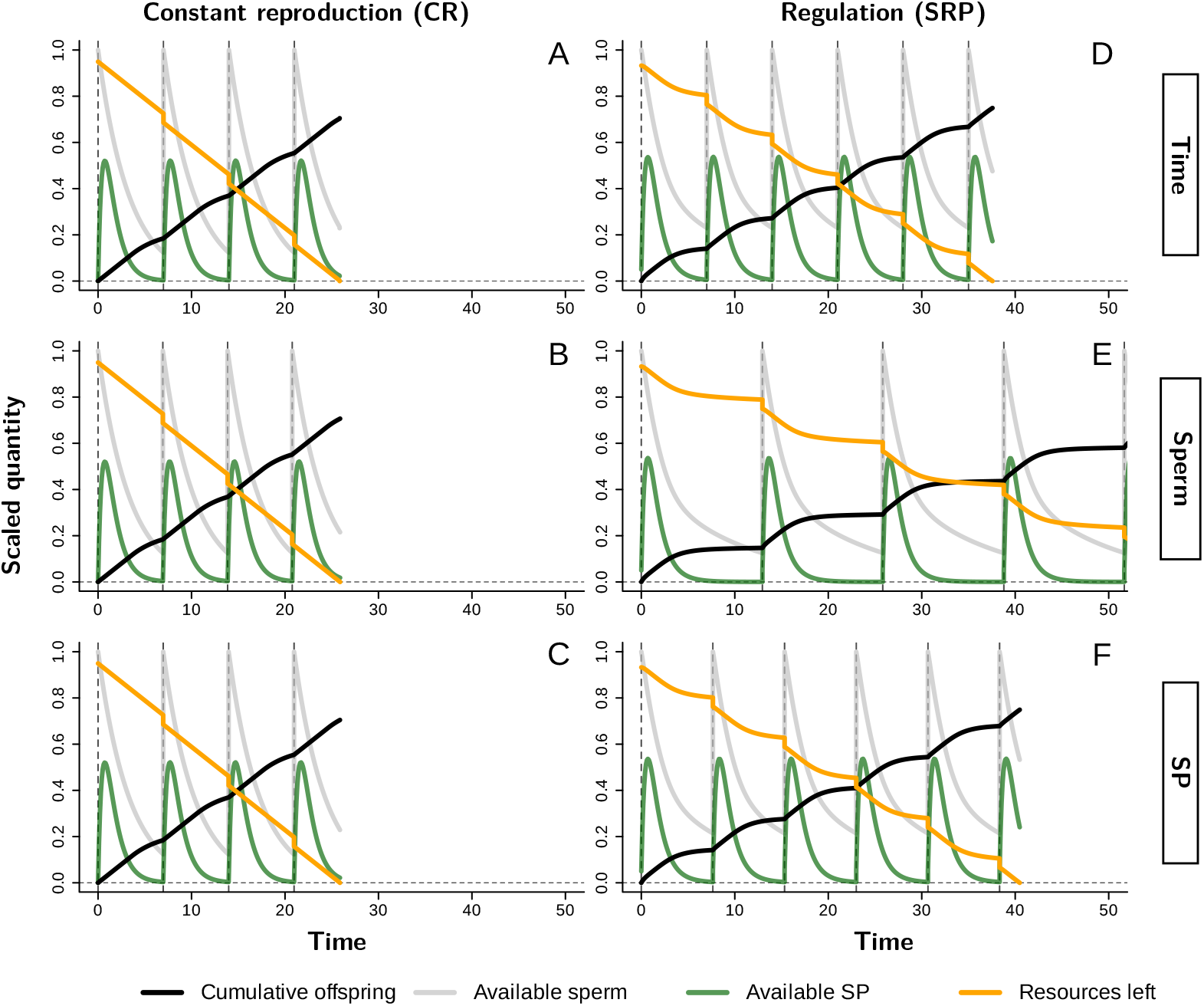
Reproductive dynamics under constant reproduction (CR) or regulation (SRP), under different stimuli of remating. To allow for representation on shared axis, all values were normalized so that maximal value (offspring and resources = 2500, sperm= *S*_0_, SP= 200) is 1. (A–C) Dynamics under constant reproduction. (A) Remating intervals are determined by the time since last mating. Female begins reproduction at time 0 when she receives sperm (gray line) and SP (green line). She produces offspring at a constant rate, using resources (orange line) to produce eggs. When the sperm becomes limiting, offspring production (black line) slows down. Female remates (dashed vertical lines) after a set time has elapsed since mating. She replenishes sperm and SP, and pays the corresponding resource cost of mating. The offspring production speeds up to its initial level, and the process repeats until resources are fully depleted. (B) Remating is determined by the number of sperm remaining in storage. Here, sperm threshold is fixed at *s* = 100, which leads to similar interval between matings as that in panel A. (C) Remating is determined by the quantity of circulating SP. It is determined by a threshold (*p* = 0.6675) which has been set to mimic the remating interval in A. In the CR scenario, SP determines only time of remating, and has no impact on egg production, release, or sperm release. (D–F) Dynamics of reproduction under SRP. (D) Time as a stimulus, as in A. Under SRP the female spends fewer resources because regulation reduces the number of unfertilized eggs she lays. This allows for more matings and a extends lifespan. (E) Sperm as a stimulus, as in B (*s* = 100). Under SRP the remating threshold is reached later, extending the interval between matings and allowing for more matings and a longer lifespan. The lifespan extends so that resources have not beenused within the plotted time. (F) SP as a stimulus, as in C (*p* = 0.6675). Again regulation extends lifespan and the interval between matings.

Reproduction under SRP changes the dynamics of offspring production—both in the patterns of sperm and egg use, and in the length of the remating interval (Fig. 2D–F). Under the three remating stimuli, offspring production shows a sharp initial acceleration which is driven by the free ovulin and SP delivered during mating, both of which trigger the release of eggs accumulated before mating. After mating offspring production remains high, sustained by the SP freed from stored sperm, which keeps stimulating egg release and production. As SP depletes and its effects wane, the egg production starts to slow down. This saves resources (*u*) by preventing the release of unfertilized eggs. It also allows females to extend their reproductive lifespan (and mate more) in comparison to the CR scenario. The dynamics under the three stimuli, identical under CR, are now different. This means that the three modeled stimuli are not affected in the same way by seminal fluid proteins. In particular, when sperm number is used as a remating stimulus (Fig. 2E), the time required to reach the necessary threshold to remate is delayed. This is caused by sperm being retained when seminal proteins are depleted, which extends the remating interval.

SRP delays resource depletion, which allows for more matings and longer remating intervals. This, in turn, increases the reproductive output: under CR female mates four times and produces 1761 offspring (Fig. 2A). Under SRP she mates six times and produces 1872 offspring (Fig. 2D). We quantified the benefits of SRP across different values of the remating stimuli (Fig. 3). To make sure that our results are robust, we also performed stochastic simulations with the initial resource budget and lifespan randomly drawn from a log-normal distribution with

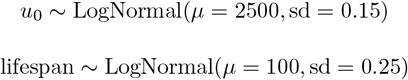

**Figure 3:**
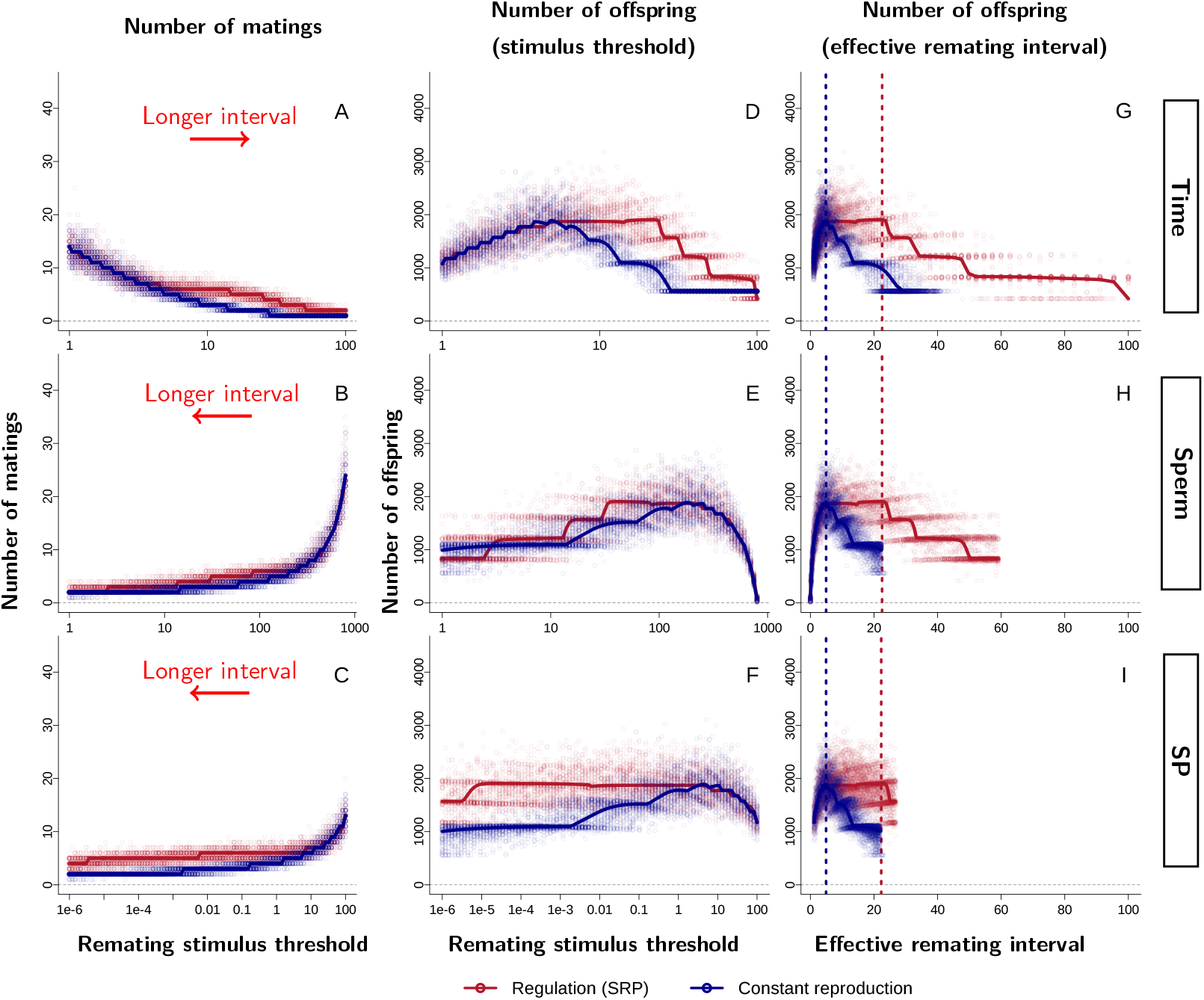
Number of matings and offspring production under the three remating stimuli. The solid lines come from the deterministic model; points represent a corresponding stochastic simulation (n=50 per each stimulus value). (A–C) Number of matings at different thresholds. Short time (A) or a high sperm/SP threshold (B–C) lead to frequent mating. Arrow shows the direction of the increasing interval between matings. There exists an intermediate remating interval where the number of matings is higher under SRP than CR—red line above the blue (e.g. panel A, threshold=10). (D–F) Offspring production at different thresholds. When remating very frequently, the mating costs consume a large fraction of the resource budget, resulting in low offspring production. The offspring production then increases as time between matings increases (D) or sperm/SP threshold decreases (E, F). However, further increase in time or decrease in thresholds leads to very long remating intervals, which results in sperm limitation— and a subsequent decline in offspring production. This manifests as a peak offspring production at an intermediate thresholds under CR. However, under SRP there is a plateau of high fitness, rather than a peak, suggesting that SRP delays the period of sperm limitation. (G–I) Offspring production as a function of the effective remating interval. Each value of the stimulus results in a specific remating interval—an *effective remating interval* which can be computed from the deterministic model. Plotting reproductive output as a function of the effective remating interval, as shown in figures G–I, allows for direct comparison across the three stimuli. The relationship between offspring production and effective remating interval is consistent across all stimuli: under CR, reproductive output is maximized at ≈5 (vertical dashed line) for all stimuli. Under SRP, fitness remains near its peak over a range of 5 to 20 days with no qualitative difference among the stimuli.

When the values of the mating stimuli are rapidly reached (i.e. high sperm/SP concentration thresholds or short time interval) time between each mating is short which leads to more matings (Fig. 3A–C). When females mate frequently (Fig. 3A, left side, Fig. 3B–C right side), the offspring production under CR and SRP is virtually the same (Fig. 3D–F). This is because fast remating allows for a frequent resupply of sperm and seminal proteins, which keeps the reproduction close to its maximum. If thresholds are reached later, offspring production increases with decrease of remating frequency. This is likely due to the decreased burden of mating costs on the total resource budget. Saved resources can be allocated to the production of more offspring. However, if thresholds cause the intervals between matings to extend so that females experience sperm limitation, offspring production decreases. This is because females lay unfertilized eggs, wasting some of their resources. Consequently, there is an optimal threshold value, leading to a not too short or too long interval between matings, at which female achieves maximum fitness.

The direct comparison of the optimal strategies under different stimuli is difficult, as the thresholds are expressed in different units (time, sperm number or SP concentration). However, each remating stimulus threshold corresponds to a time interval between matings, which we can compute from our deterministic model. In practice, we averaged time periods between matings to obtain this interval, referred to as the “effective remating interval”. This interval excludes the period from the last mating to the end of reproduction, as this last period is determined by resource or lifetime constraints and thus does not reflect a remating strategy. When expressed in terms of effective remating intervals (Fig. 3G–I), the three stimuli point to the same optimal remating strategy. The only observable difference between the stimuli is that very long intervals are never reached when females use sperm or SP as cues of remating (Fig. 3H–I). This indicates that the long intervals would require thresholds lower than what we considered biologically realistic.

Figures 3G–I further demonstrate how SRP changes the optimal remating strategy. In the absence of any regulation (CR model), offspring production reaches a peak at an intermediate interval of approximately five days, immediately followed by a sharp decrease. The decrease comes from egg production being kept high even when sperm are limiting. The female then produces unfertilized eggs and wastes resources, which in turn reduces the possibility to remate and to produce more offspring. Conversely, with regulation (SRP model), egg production is down-regulated when sperm are limiting. The saved resources allow for additional matings, resulting in a situation where the number of matings is higher under SRP than CR (e.g. figure 3A, for a time interval of 10). As a consequence, under SRP (with parameters set as in the table of Fig. 1B), offspring production stays close to its maximum from approximately five to 20 days (which actually yields maximum offspring production in this case). The regulation of female reproductive physiology by SP therefore allows females to maintain high offspring production over a wide range of time intervals between matings.

Under the assumptions of our SRP model, the number of offspring is close to maximum over a range of possible remating intervals. This raises the question of whether the shift from peak-like maximum, under CR, to a broader plateau under SRP occurs abruptly or gradually, as seminal proteins increasingly influence the regulation of female reproduction. To answer this question, we varied the degree of influence of seminal fluid proteins on female physiology. We defined the degree of regulation as the difference between the base and the maximum values of the three quantities which are sensitive to SP (egg production, egg release and sperm release). A degree of regulation of 0 means that the base and maximum values of those quantities are equal so that seminal proteins have no influence on these processes—equivalent to the CR scenario. A degree of regulation of 1 corresponds to the SRP scenario (values as in Fig. 1B). Intermediate degrees of regulation are modeled by keeping the maximum values constant and adjusting the base values. For example, egg production under the SRP scenario (regulation degree 1) can take values from *δ*_0_ = 15 to *δ*_max_ = 120 (Fig. 1B). A regulation degree of 0.2 is obtained by increasing the base value *δ*_0_ to 120 − 0.2(120 − 15) = 99 while keeping *δ*_max_ constant.

We first explored the effect of regulation degree under the same conditions as in previous simulations— specifically, with the cost of mating fixed at 100. Time was used as the stimulus, because our earlier results showed that the nature of the stimulus does not affect the optimal remating strategy (Fig. 3). These simulations demonstrate that the offspring production is consistently maximized at an intermediate remating intervals (Fig. 4A). However, as the degree of regulation increases the optimal strategy shifts toward longer intervals, and the range of intervals over which reproductive output remains near its maximum expands. Thus regulation creates a sort of buffer that allows females to produce many offspring even when the optimal remating interval cannot be achieved. This pattern appears independent of mating cost: increasing mating cost to 500, as expected, reduces the number of produced offspring but does not change how regulation shapes the relationship between offspring production and remating interval (Fig. 4B).

**Figure 4:**
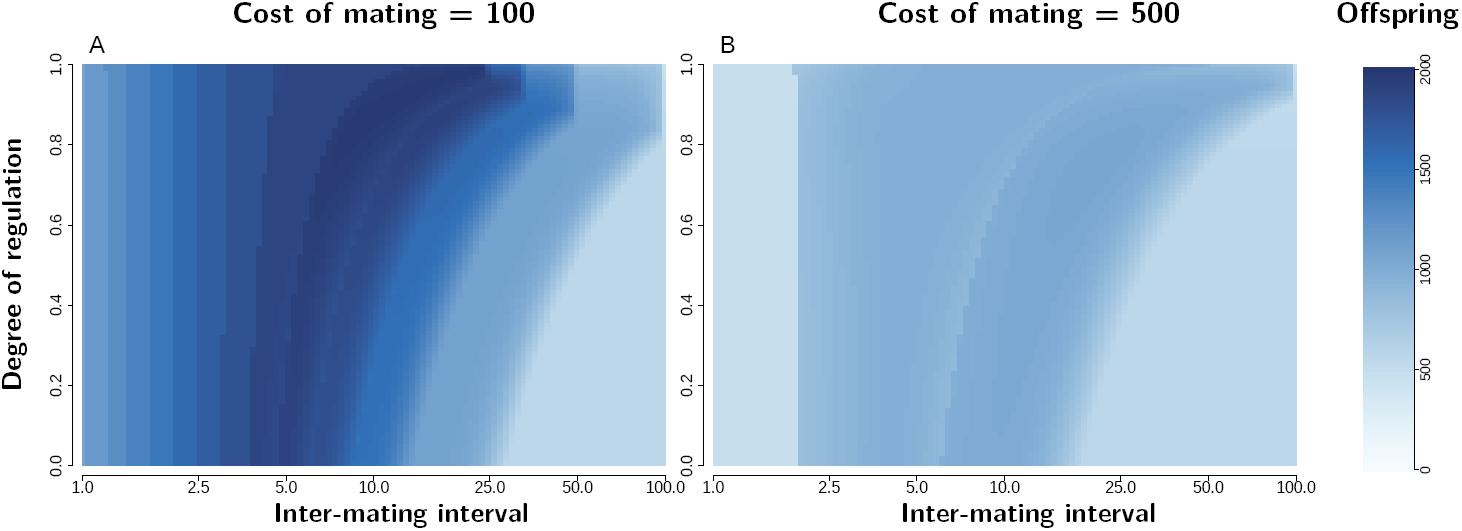
Offspring production as a function of the interval between matings for varying degree of regulation and different mating costs. In these simulations, remating is triggered by time. The degree of regulation describes the influence that SP and ovulin have on the female reproduction with 0 corresponding to the CR scenario and 1 to the SRP scenario. (A) Cost of mating is 100, as in previous simulations. For any degree of regulation, offspring production is maximized at an intermediate remating interval. When the degree of regulation increases, the optimal strategy shifts to a longer interval and range of high offspring production is extended. (B) Increasing mating cost to 500 reduces the number of offspring, but does not change the way regulation affects offspring production.

## 4 Regulation by SP-like seminal proteins can reduce intersexual conflict

Previous analysis demonstrates that females maximize their offspring production at intermediate remating intervals, and that regulation by seminal fluid proteins keeps offspring production maximum when remating is delayed. Delayed remating should be in males’ interest, as it will likely increase their contribution to offspring produced by the female. However, despite the delay, female optimal interval can still be shorter than that of males. In other words, there remains a possibility of an intersexual conflict over the remating time.

To investigate how seminal fluid proteins influence conflict over remating, we modeled a scenario in which males can manipulate females’ remating interval. We first considered a population in which all males impose an effective remating interval that aligns with the females’ optimum. We then considered mutant males which impose a different remating interval. Because mutants are rare, each female encounter at most one mutant during her lifetime. We calculated the fitness of mutant males by computing their offspring production at each possible position within a female mating sequence and averaging across positions, assuming all positions are equally probable. For simplicity, we assumed that the sperm of a mutant male entirely displaces that of previous mates and is itself entirely displaced by the sperm of subsequent mates. Proceeding this way, we determined which strategy is optimal for mutant males. When this strategy does not align with that of the female, it means sexual conflict occurs.

Mutant males consistently benefit from longer remating intervals under both CR and SRP scenarios (Fig. 5A). This conclusion holds even when non mutant males induce a remating interval that exceeds females’ optimum (e.g. 30 time units in Fig. 5B). In such cases, mutant males that impose an even longer intervals are able to invade the male population. Together, these simulations predict a clear sexual conflict over remating, as predicted by theory, with selection always favoring males that impose the longest remating time.

**Figure 5:**
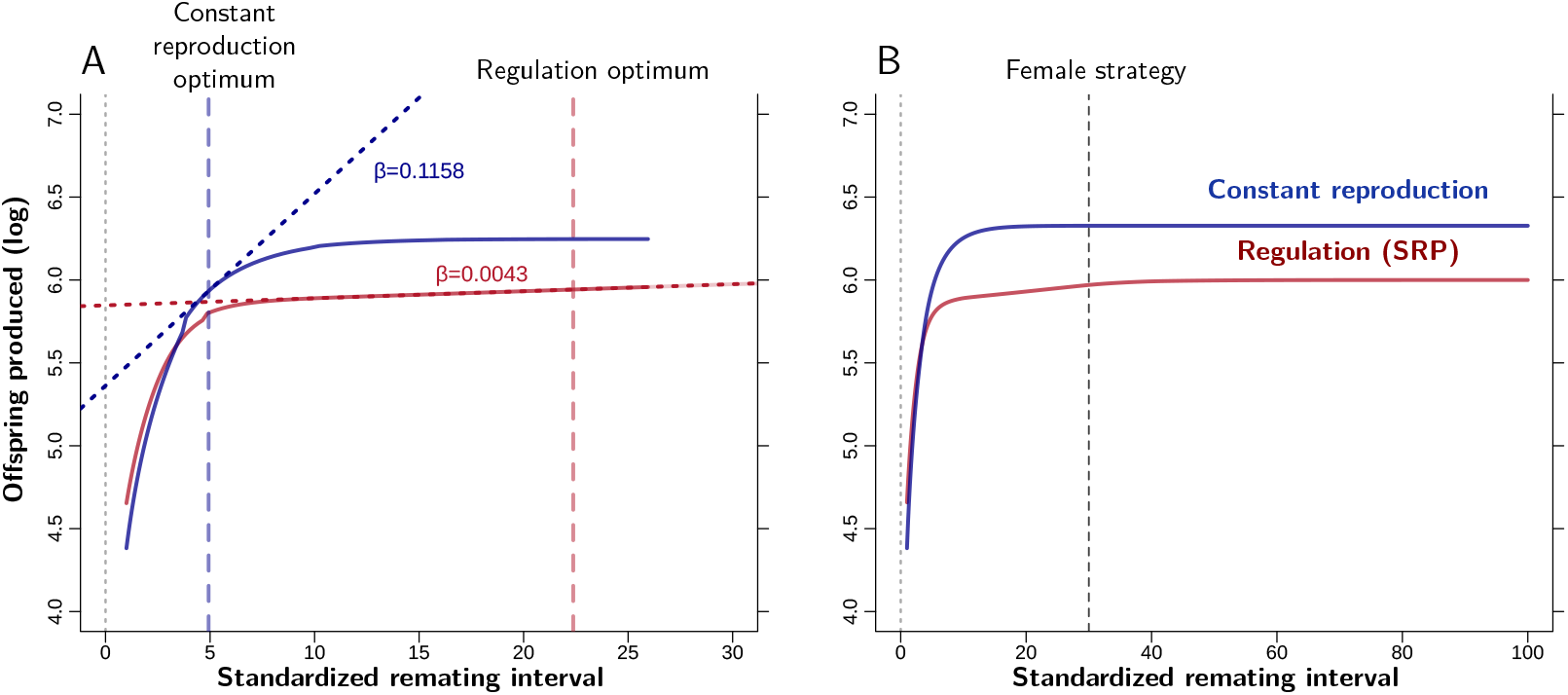
**(A) Regulation by SP-like proteins reduces sexual conflict over remating interval.** We computed the fitness of mutant males that manipulate female remating behavior in a context where non-mutant males allow females to follow their optimal remating strategy. Male fitness increases monotonously with remating interval, indicating a sexual conflict over remating. We quantified the strength of this conflict as the slope of the relationship between male fitness (log scale) and remating time, evaluated at female optimum (see text for further details). In the absence of regulation (CR), the selection gradient is 0.115. With regulation (SRP), it drops to 0.004, indicating a substantially weaker conflict between the sexes. **(B) Mutant’s fitness when the majority of males enforce a long intermating interval** Even when the majority of males imposes an interval that greatly exceeds female optimum (here 30 days) mutants that impose even longer intervals have higher fitness. By contrast, mutant imposing a shorter interval cannot successfully invade the population.

However, although this conflict appears unavoidable, its strength may vary with the degree of regulation. We quantified the strength of sexual conflict as the fitness benefit males gain by increasing the interval between matings beyond the female optimum. Specifically, we measured the slope of the relationship between the mutant male log fitness and the remating interval they impose, evaluated at female optimum. This slope corresponds to the strength of selection acting on males, as defined by the selection gradient *β* (47). A steeper slope indicates stronger selection and, correspondingly, a more intense sexual conflict.

We found that the slope under CR is steeper than that under SRP (*β* = 0.116 vs *β* = 0.004, see Fig. 5A), indicating that selection on males to extend female optimum is weaker under SRP. Thus, regulation of female reproduction by seminal fluid proteins mitigates the sexual conflict over remating, and may render it negligible relative to other selective forces.

We further investigated this mitigation of the sexual conflict by computing females’ optimal remating strategy and the corresponding gradient of selection under varying degree of regulation and across different mating costs (Fig. 6). We only present computation performed with time as a stimulus, as the three stimuli are functionally equivalent (Fig. 3). Increasing the degree of regulation leads to longer intervals between matings (Fig. 6A). As the interval extends, female mates fewer times (Fig. 6B), which generally leads to a decreasing selection gradient and thus a weaker sexual conflict (Fig. 6C).

**Figure 6:**
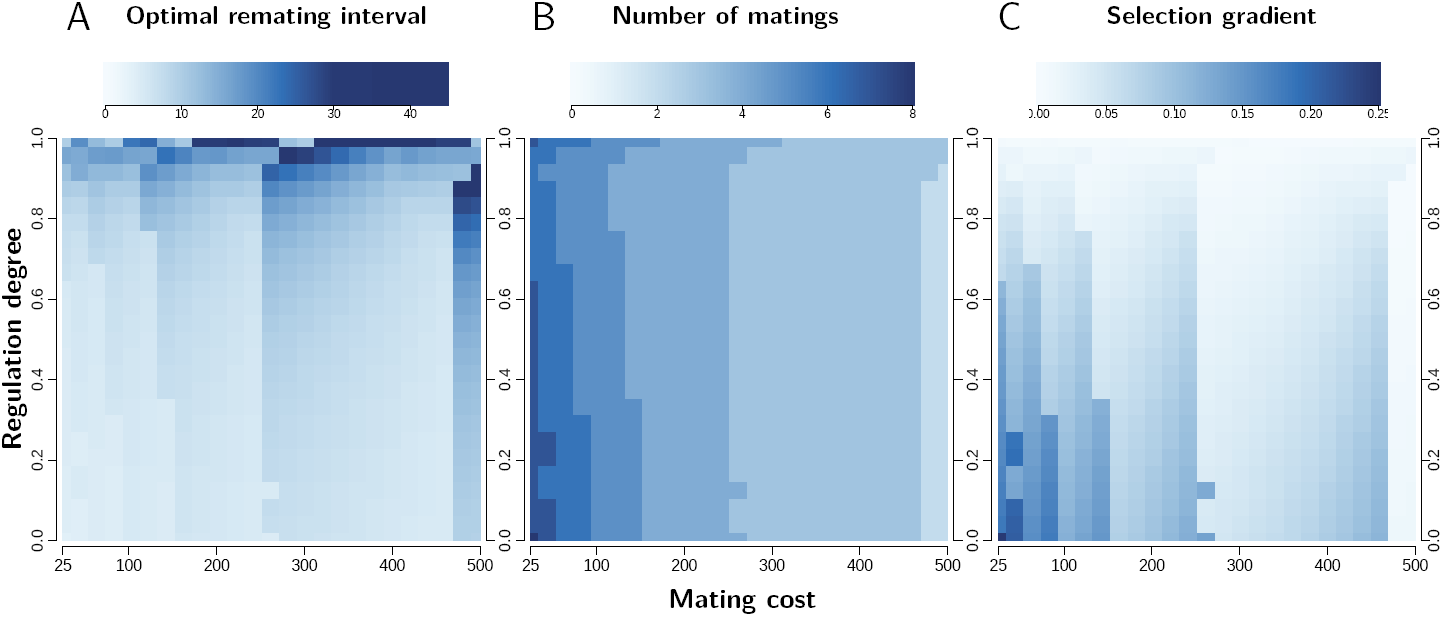
The conflict over remating as a function of regulation degree and mating costs. Each tile represents a calculation of optimal interval and of strength of sexual conflict, similar to those in figure 5 for varying regulation degree and mating cost. (A) Females’ optimal remating interval as a function of mating cost and regulation degree. Increasing the cost of mating or the degree of regulation leads to longer optimal remating interval. (B) The number of matings per female follows the inverse trend. Females mate only a few times when the cost of mating or the degree of regulation are high. (C) The resulting selection gradient, used here as a measure of sexual conflict over remating interval, generally decreases with decreased number of matings. As a result, the sexual conflict decreases with both mating cost and the increased degree of regulation. The discontinuities result from the discrete number of matings.

Varying the cost of mating alters the intensity of sexual conflict, though not in a straightforward manner. In principle, increasing mating costs reduces the benefits of additional matings for females and should therefore promote greater alignment between male and female interests. This explains the stepwise decrease in conflict observed in Fig. 6B, with each step corresponding to a reduction of one mating event in the female optimal strategy. A less expected feature is that when mating costs increase, yet not enough to reduce the number of matings (e.g. from 300 to 450 in Fig. 6B), the female optimal remating interval actually decreases, thereby strengthening the sexual conflict.

This pattern arises because sperm depletion causes offspring production to decline faster than linearly with time. As a result, total reproductive output is maximized when a female spends equal time utilizing each mate’s sperm. Equivalently, a female that mates *k* times over a total period *T* has an optimal remating interval of *T/k*. If mating costs increase while the number of matings remains constant, *T* decreases due to reduced resources for offspring production, and the optimal remating interval decreases accordingly. By contrast, a small increase in mating costs that reduces the number of matings from *k* to *k* − 1 increases the resources available for reproduction, thereby increasing *T*. The combination of fewer matings and a longer reproductive period then leads to a longer optimal remating interval.

## 5 Discussion

Our model explores how seminal fluid proteins regulate female reproduction, based on the described effects of sex peptide (SP) (reviewed in 3) and ovulin. In our previous work, we showed that seminal fluid proteins could provide a timed signal to decrease egg production when sperm become limiting (38). The resulting alignment of sperm use and egg production improved reproductive efficiency, highlighting both the general importance of this alignment, and the role of seminal fluid proteins in determining female fitness.

Here, we extended this analysis to multiple matings and the downstream effects of regulation on remating dynamics. In agreement with the prevailing theory, we found that males and females are in conflict over the time of remating: while males benefit from the longest possible time, females have an intermediate optimal remating time. However, we found that the intensity of this conflict is attenuated by the effects of SP on females’ reproductive physiology.

The attenuation of sexual conflict comes from females’ optimal remating time being longer under SP regulation than under Constant Reproduction. This is possible because the concentration of SP-like proteins is a reliable indicator of the remaining sperm in storage (38), which females can use to coordinate the release of sperm from storage with egg availability. In the model, this coordination reduces the wastage of energy at longer intervals.

The coordination hypothesis finds a good empirical support in *Drosophila melanogaster*, where SP might be an effective cue for monitoring the number of sperm left in storage. SP is initially attached to sperm, and must detach before being active, making the rate of SP depletion reflects that of sperm release. In addition, the neurons detecting SP are located both near the spermathecae (the sperm storage location) and near the uterus exit, making it a good candidate for the simultaneous control of sperm and egg release (48). These factors are also consistent with empirical observations that the probability of remating correlates with sperm number in storage (49). Beyond the specific case of *Drosophila*, other species have similar regulatory mechanisms, even when the regulation is not directly governed by seminal fluid proteins. For example, in mosquitoes, prolonged release of 20-hydroxyecdysone from the digestion of the mating plug might provide an analogous cue that females could use to decrease egg production over time (50). Our results indicate that this type of signal plays a pivotal role in mitigating sexual conflict, yet remains largely overlooked and insufficiently investigated.

We predict that sexual conflict over remating should be strongest in the absence of SP regulation (but for contrasting experimental results see (51)). This is based in part on the simplifying assumption that the number of offspring produced by a male depends only on how long the female utilizes his sperm. However, this number might also vary with patterns of sperm competition. In our model, we assumed total sperm displacement at each new mating. This simple modality of sperm competition generates an extreme form of sexual conflict, as female remating results in the complete loss of any remaining potential reproductive success for the initial male. In principle, partial displacement would simply lower the conflict, without altering our quantitative conclusions. More complex situations could arise in case of preferential use of sperm from earlier males, which would create asymmetry between males mating at different times, or if SP and other seminal fluid proteins play an active role in modulating sperm competition. This dual capacity of seminal fluid proteins to regulate female physiology and modulate sperm competition has been demonstrated in several instances (52–57). Its consequences requires further exploration, but exceeds the scope of this paper.

We have also assumed in our model that females have a fixed resource budget, set at the beginning of their reproductive life, encapsulating reproduction-lifespan trade-off. In many species however, mating is associated with extra resources because males provide nutrients in the ejaculate, or nuptial gifts that females eat (36, 58–60). We did not explicitly address this situation, which might reduce the benefits coming from saved energy from decreasing egg production. Although letting females acquire additional resource when remating is largely equivalent to reducing the cost of mating in our model, females could also change their resource intake dynamically, and nutritional availability itself can influence the pattern of egg laying (58, 61). This could in turn affect egg production, and change the energy balance. It is also known that in *Drosophila*, SP itself can increase both resources expenditure and intake (10, 29, 62). This means that the patterns of energy use and mating cost are probably not the same between CR and SP regulated reproduction, the meaning of which might be understated by the simplified model of resource consumption in the model.

However, the proposed model generates predictions that extend beyond the limitation of sexual conflict over remating. Compared to the situation with no regulation, where their reproductive output peaks at an intermediate time value, females who regulate their reproduction based on SP have a close to maximum reproductive output across a range of different intervals. This could provide females with a buffer and allow flexibility of the remating interval when mating is temporarily impossible, due to lack of partners or unfavorable environment, like lack of oviposition sites or dangers of predation. Such buffer would also allow for increased selectivity, allowing females to wait until they encounter a high-quality male to remate with. The selectivity hypothesis has some empirical support in *Drosophila*, where mated females show a decreased sensitivity to male pheromones, resulting in higher selectivity, which extends remating intervals (28, 63). The physiological mechanism involves the juvenile hormone, which dynamics is tightly connected to mating. In many insect species (18, 64– 66), juvenile hormone is indeed either transferred directly in seminal fluid, or its production by females is upregulated by seminal fluid proteins (including by SP). Increased level of juvenile hormone raises the pheromone threshold required to stimulate female choice (28, 67), leading females to prefer higher-quality males (63). The same reduction in female sensitivity is also observed in Tephritidae, where mated females show reduced attraction to male pheromones compared to virgins (68). Our model shows how the regulation of female reproduction by SP, and the efficient egg use it permits, could decrease the fitness cost of such male selectivity.

Taken together, the results from our model frame seminal fluid proteins as a valuable source of information. They shift the focus from males manipulation of female behavior toward females utilizing seminal fluid proteins to regulate their physiology, as part of their own reproductive strategy. It might seem counter-intuitive that males have such limited control over the action of the proteins, which they produce and provide to females. However, there is empirical evidence that females do in fact have an active role in the utilization of the seminal fluid proteins. In *Drosophila melanogaster*, SP can fully bind to sperm only once inside the female reproductive tract, giving females control over its activation (69, 70). And in the potato weevil *Euscepes postfasciatus*, while females do require seminal fluid proteins for the post-mating refractory period to occur, the length of this period depends largely on the female, remaining the same across different males (71, 72). In addition, if males and females were in conflict over the utilization of seminal fluid proteins, we would expect male proteins to co-evolve with female receptors in an evolutionary arms race. However, there is no evidence of such a coevolution in *Drosophila* between SP and its receptor. The fragment of SP responsible for stimulating fecundity is highly conserved, and the sex-peptide receptor appears to have evolved under selective pressures reflecting constraints from functions beyond that of binding to SP (73).

In this paper, we have presented a formal exploration of how physiological adjustments of egg and sperm use, caused by seminal fluid proteins, affect optimal remating intervals. Our model represents a scenario in which seminal fluid proteins are the primary regulatory mechanism, allowing us to distill the potential role of physiological regulation by these proteins. As females modify their mating rate based on the sperm availability and, given that seminal fluid proteins increase egg production and sperm usage, it is likely that seminal fluid proteins change the pattern of remating as a downstream consequence. The results paint the picture of seminal fluid proteins as largely beneficial to females. They reduce female resource wastage and permit greater mate selectivity. And, although experimental data shows that factors such as nutrition and sperm competition patterns could influence these results (51, 74), we find that seminal fluid proteins can reduce sexual conflict over the time of mating. Under the assumptions of the model, the broad physiological changes caused by seminal fluid proteins are unlikely to be purely antagonistic and the results question the primary role of sexual conflict behind the evolution of seminal fluid proteins.

## AUTHORS CONTRIBUTION

P.M., D.D. and J.B.F. conceptualized the study; P.M. and J.B.F. coded the model in C++; P.M. analyzed the results and led the writing of the manuscript with assistance from D.D. and J.B.F.; All authors contributed critically to the drafts and gave their final approval for publication.

## DECLARATION OF COMPETING INTERESTS

Authors declare no competing interests.

## ACKNOWLEDGMENTS

We thank Thomas Flatt and Sébastien Lion for thoughtful discussions throughout the project, and Mariana Wolfner, Hanna Kokko and Jennifer Perry for their valuable comments on the manuscript. P.M. was supported by PhD funding from Université Toulouse 3 – Paul Sabatier through the SEVAB doctoral school. D.D. was supported by a FCT fellowship (2023.08149.CEECIND).

## Notes

### Competing Interest Statement

The authors have declared no competing interest.

